# Neuronal and astrocytic adaptations in the lateral habenula during withdrawal from chronic ethanol

**DOI:** 10.64898/2026.08.04.742855

**Authors:** Karl Y Bosque-Cordero, Shikun Hou, Elizabeth J Glover

## Abstract

The lateral habenula (LHb) encodes aversive states and negative affect, positioning it as a candidate region for the negative reinforcement that drives alcohol withdrawal. However, little is known about how chronic ethanol exposure affects LHb neuronal function and glial biology during withdrawal. Here, we used chronic intermittent ethanol (CIE) vapor exposure, a well-established model of alcohol dependence that reliably produces somatic and affective signs of withdrawal, to examine LHb physiology and astrocytic markers during acute withdrawal in male and female rats. Whole-cell and cell-attached recordings revealed that withdrawal reduced evoked and spontaneous firing in LHb neurons, with rebound firing following a crossover pattern between males and females. Despite these excitability changes, the overall distribution of firing phenotypes was unchanged, suggesting a shift in gain rather than a reorganization of cell types. Immunofluorescence revealed increased Sox9+ and GFAP labeling in the LHb during withdrawal at the same time point when electrophysiology experiments uncovered impaired astrocytic regulation of glutamate clearance. Together, these findings reveal that withdrawal from chronic ethanol exposure produces neuronal and glial adaptations in the LHb, pointing to impaired glutamate regulation as a candidate mechanism relevant to the negative affective state of alcohol withdrawal. These findings position the LHb as a potential node linking astrocyte-neuron dynamics to withdrawal symptoms and relapse vulnerability in alcohol use disorder.

## Introduction

Alcohol use disorder (AUD) is a significant public health concern characterized by compulsive alcohol consumption and impaired control over intake (American Psychiatric Association, 2013). In the United States alone, an estimated 27.9 million individuals aged 12 and older, or 9.7% of that population, met diagnostic criteria for AUD in 2024 (SAMHSA, CBHSQ., 2025). Among those attempting to abstain from alcohol, roughly half experience a constellation of withdrawal symptoms including anxiety, insomnia, hyperalgesia, irritability, and anhedonia that frequently contribute to relapse (American Psychiatric Association, 2013). Despite this, there are currently no pharmacotherapeutics with FDA approval for the treatment of withdrawal symptoms in patients with AUD (Breese et al., 2005; Kushner et al., 2000; Schuckit, 2014). This gap is particularly consequential because withdrawal symptoms increase relapse risk two-to three-fold, with severe cases showing relapse rates of 70 to 80% within three months (Booth & Blow, 1993; Miller, 2001). These data underscore the urgent need to better understand the neural mechanisms that drive withdrawal symptomology in order to uncover new pharmacotherapeutic targets that can disrupt the withdrawal-relapse cycle that perpetuates AUD.

The lateral habenula (LHb) has emerged as a critical brain region involved in encoding aversive signals, regulating affective states, and modulating pain sensitivity. As a predominantly glutamatergic nucleus, the LHb serves as a major output hub, projecting to midbrain structures including the rostromedial tegmental nucleus (RMTg), which in turn inhibits ventral tegmental area (VTA) dopamine neurons (Jhou, Fields, et al., 2009; Jhou, Geisler, et al., 2009; Stamatakis et al., 2013). Through this disynaptic circuit, the LHb plays a pivotal role in negative reward prediction, suppression of reward signaling, and encoding of nociceptive and anxiogenic states (C. Chen et al., 2024; T. Chen et al., 2024; X. Li et al., 2024) positioning it as a leading candidate neural substrate in which withdrawal symptoms may manifest.

With this in mind, a number of studies using voluntary drinking models have examined the effects of ethanol on the LHb, establishing it as a site of ethanol-induced neuroadaptation and implicating it in the behavioral changes associated with ethanol exposure (Flanigan et al., 2023; Haack et al., 2014; Kang et al., 2017; J. Li et al., 2016). However, the effects of ethanol dependence on LHb physiology are still not understood. Importantly, LHb neurons display highly heterogeneous firing patterns—silent, tonic regular, tonic irregular, and bursting—that are thought to reflect underlying differences in intrinsic membrane properties, synaptic input strength, and channel conductance profiles (Wagner et al., 2017; Weiss & Veh, 2011; Wilcox et al., 1988). Moreover, distinct LHb firing phenotypes have been linked to pathological states (Cui et al., 2018, 2019; Fan et al., 2023; Ma et al., 2024; Y. Yang et al., 2018). Yet, how chronic ethanol exposure affects phenotypically distinct LHb neurons or if it destabilizes the balance across firing phenotypes has not been explored.

The present study was designed to address this gap by measuring changes in excitability and synaptic transmission in phenotypically distinct LHb neurons following chronic ethanol exposure in a model of dependence that is well-characterized for producing robust physical and affective symptoms of withdrawal (Gilpin et al., 2008, 2009; Glover et al., 2019; Mitten et al., 2026; Nentwig et al., 2022; Trantham-Davidson et al., 2014). Our findings reveal that single LHb neurons exhibit stochastic transitions among distinct firing phenotypes that are unaffected despite robust changes in LHb excitability and astrocytic regulation of synaptic glutamate during acute withdrawal from chronic ethanol exposure.

## Methods

### Animals

Adult male and female Long-Evans rats (P60 at arrival; Envigo, Indianapolis, IN) were used in all experiments. Rats were allowed to acclimate to the facility for at least seven days prior to the initiation of any experimental procedures. Rats were singly housed in standard polycarbonate cages in a temperature-controlled vivarium maintained on a reverse 12:12 h light/dark cycle (lights on at 22:00). Rats had *ad libitum* access to standard chow (Teklad 7912, Envigo) and water throughout the study. All procedures were approved by the University of Illinois Chicago Institutional Animal Care and Use Committee and conducted in accordance with the National Institutes of Health (NIH) Guide for the Care and Use of Laboratory Animals.

### Chronic intermittent ethanol vapor exposure

Rats were rendered dependent using a chronic intermittent ethanol (CIE) vapor exposure paradigm that is well-characterized for producing both somatic and affective signs of withdrawal (Gilpin et al., 2008, 2009; Glover et al., 2019; Roberts et al., 1996, 2000). Rats that were assigned to the CIE group were exposed to vaporized ethanol in custom-built chambers for 14 hours/day (18:00–08:00) over a 14-day period. Control rats (AIR group) were exposed to ambient room air under identical conditions. Behavioral signs of intoxication were scored daily in CIE-exposed rats using a five-point subjective rating scale (1 = no signs of intoxication; 5 = complete loss of consciousness), as described previously (Glover et al., 2019, 2021; Ramirez et al., 2024).

Blood samples (40 μL) were collected via tail nick from CIE-exposed rats immediately after vapor exposure on days 2, 7, 10, and 14 (±1 day). Controls received a tail-pinch at matched timepoints to approximate the level of discomfort experienced by intoxicated rats during blood sampling. Blood samples were centrifuged at 10,000×g for 10 minutes at 4 °C, and the plasma supernatant (20 μL) was aliquoted into sterile 0.5 mL microcentrifuge tubes and stored at −20 °C. Blood ethanol concentration (BEC) was quantified using an Analox Alcohol Analyzer (Analox Instruments Ltd., UK).

### Patch-clamp slice electrophysiology

Brain slices containing the LHb were prepared for patch-clamp slice electrophysiology using previously published procedures (Glover et al., 2023; Przybysz et al., 2024). Twenty-four hours following the final vapor exposure session, rats were deeply anesthetized with isoflurane and rapidly decapitated. Brains were quickly extracted and immersed in ice-cold, oxygenated cutting ACSF composed of (in mM): 125 NaCl, 2.5 KCl, 1.25 NaH₂PO₄, 25 NaHCO₃, 10 glucose, 0.4 ascorbic acid, 4 MgCl₂, and 1 CaCl₂. Coronal slices (220 μm) containing the LHb were prepared using a vibratome (Leica Microsystems) and incubated at 34 °C in oxygenated ACSF (95% O₂/5% CO₂) for at least 1 hour prior to recordings. For recordings, slices were transferred to a submerged recording chamber perfused with oxygenated ACSF at 1 mL/min and maintained at 34 °C. Recording ACSF had the following composition (in mM): 125 NaCl, 2.5 KCl, 25 NaHCO₃, 10 glucose, 0.4 ascorbic acid, 1.3 MgCl₂, and 2 CaCl₂, pH adjusted to 7.2–7.4, with osmolarity ∼300 mOsm. All recordings were conducted from 1 to 4 hours after slices were removed from the incubation chamber and were performed using an Axon Multiclamp 700B amplifier and Digidata 1550A digitizer (Molecular Devices), controlled by pClamp 11 software running on Windows 10. Neurons were visualized via infrared differential interference contrast (IR-DIC) microscopy (Olympus America). Patch pipettes (3–5 MΩ resistance) were pulled from thin-walled borosilicate glass (Warner Instruments) using a P-97 puller (Sutter Instruments). Series resistance and membrane properties were monitored throughout recordings; cells with >15% change in access resistance were excluded from analysis. Liquid junction potentials were not corrected. Data were analyzed offline using pCLAMP 11 software.

### Whole-cell current clamp recordings

To measure CIE-induced changes in intrinsic excitability, pipettes were filled with a potassium gluconate-based intracellular solution containing (in mM): 125 K-gluconate, 20 KCl, 10 HEPES, 1 EGTA, 2 Mg-ATP, 0.3 Na-GTP, 10 phosphocreatine, and 0.02 MgCl₂; pH adjusted to 7.3, ∼285 mOsm. LHb neurons were held at −70 mV, and 500 ms current steps (−100 to +220 pA in 20 pA increments) were applied.

### Whole-cell voltage clamp recordings

To assess spontaneous synaptic activity, pipettes were filled with a cesium methanesulfonate-based solution containing (in mM): 135 cesium methanesulfonate, 20 KCl, 1 MgCl₂·6H₂O, 0.2 EGTA, 4 Mg-ATP, 0.3 Na-GTP, 20 phosphocreatine, and 2.91 QX-314; pH adjusted to 7.3, ∼290 mOsm. Cells were first clamped at −55 mV for 5 min to record spontaneous excitatory postsynaptic currents (sEPSCs) after which they were clamped at +10 mV for 5 min to record spontaneous inhibitory postsynaptic currents (sIPSCs) according to previously published procedures (Przybysz et al., 2024). The frequency and amplitude of synaptic events were captured and excitatory/inhibitory (E/I) balance was quantified using two complementary approaches according to previously published procedures (Przybysz et al., 2024). First, the E/I frequency ratio was calculated by dividing the frequency of sEPSCs by the frequency of sIPSCs within each cell. Second, synaptic drive, which provides a more comprehensive measure of the relative strength of net excitatory versus inhibitory synaptic input by incorporating both the frequency and amplitude of synaptic events, was assessed by calculating the ratio of excitatory to inhibitory input strength, defined as the product of sEPSC frequency and amplitude divided by the absolute value of the product of sIPSC frequency and amplitude. The absolute value of the inhibitory component was used to prevent sign inversion due to the inward (excitatory) versus outward (inhibitory) directionality of postsynaptic currents.

### Spontaneous firing recordings

Spontaneous firing was measured using two configurations. First, cell-attached recordings were performed using pipettes filled with the same potassium gluconate-based intracellular solution described above. Cell-attached configuration was achieved by applying light positive pressure to the pipette until contact was made with the neuronal membrane, at which point gentle suction was used to form a high-resistance seal (>1 GΩ). No additional suction was applied, and the membrane was not ruptured, thereby preserving the intracellular milieu. Neurons were held at pipette potential (0 mV command voltage), and spontaneous action potential-associated capacitive currents were recorded for 3 minutes.

Immediately following the 3-minute cell-attached recording, the membrane patch was ruptured to establish whole-cell configuration, and spontaneous firing was recorded for an additional 3 minutes in current-clamp mode in the absence of any current injection (I = 0 pA). This paired cell-attached-to-whole-cell approach allowed within-cell comparison of firing phenotype before and after membrane rupture, controlling for potential effects of intracellular dialysis on firing pattern.

### Astrocytic regulation of synaptic glutamate

To assess the contribution of astrocytic glutamate transporter activity on LHb firing, baseline spontaneous activity was recorded in a subset of neurons in cell-attached configuration as described above. This was followed by a second recording of spontaneous activity after bath application of the GLT-1 inhibitor, dihydrokainic acid (DHK; 100 μM), for 10 minutes.

### Firing phenotype classification

Spontaneous firing during cell-attached and whole-cell I=0 recordings was classified into four mutually exclusive phenotypes using a custom MATLAB (MathWorks, Natick, MA) pipeline (GitHub). Action potentials with an inter-spike interval (ISI) <1 ms were excluded as noise. To identify bursts, action potentials separated by an ISI <100 ms were first linked into candidate clusters; a cluster was confirmed as a burst only if the ISI initiating it (i.e., the interval between the last non-clustered spike and the first spike of the cluster) was ≤21 ms, distinguishing genuine high-frequency onset from loosely spaced spikes that happened to fall within the 100 ms linking window. This two-step criterion is consistent with T-type Ca²⁺ channel-mediated burst firing described in LHb neurons (Cui et al., 2018; Y. Yang et al., 2018), and previous firing classification in the LHb (Wagner et al., 2017). Absence of firing (i.e., silence) for ≥1500 ms was classified as a pause. Silence at the onset of recording, prior to the first action potential, was classified as a pause under the same 1500 ms criterion, using the start of the recording epoch as the reference boundary in place of a preceding spike. Any residual silence of <1500 ms that extending to the end of the 3-minute recording after the last detected activity segment was categorized as a “gap” rather than a pause, since it reflected the recording’s fixed endpoint rather than a true post-firing silent period. The remaining non-burst spike trains of ≥4 spikes were classified as tonic regular if the ISI coefficient of variation (CV) was ≤0.30, or tonic irregular if the ISI CV was >0.30 or if fewer than 4 spikes were present, consistent with regular/irregular tonic firing distinctions (Wagner et al., 2017).

The contribution of each phenotype to a given recording was quantified as percent of total recording duration. For every activity segment within a recording, duration was calculated as the duration of that activity segment plus its leading inter-event interval (i.e., the silent gap preceding an activity segment was credited to that activity segment), summed within each phenotype category, and divided by total recording length (%Burst, %TonicRegular, %TonicIrregular). The percent duration of time spent in post-firing pause and the trailing gap in firing that occurred prior to the end of the 3-minute recording (i.e., the time following the end of the last detected activity segment) were computed the same way. Thus, for a given recording, the percent duration for all five phenotype categories summed to 100%. Each recording was then assigned an overall phenotype corresponding to the active firing phenotype that accounted for the largest percentage of recording duration, with post-firing pause and gap excluded from this classification.

### Statistical analysis

Conventional analyses were performed using GraphPad Prism 11 (GraphPad Software, Boston, MA). Between-group comparisons were conducted using unpaired t-tests and ANOVAs with vapor group, sex, current step, or pharmacological treatment as factors where appropriate. Paired or repeated-measures approaches were used for within-subjects analyses. Posthoc comparisons were conducted using Šídák’s or Bonferroni’s correction as indicated. Because changes in the proportion of a given firing phenotype represents a change to a part of a whole, these data were analyzed using a centered log-ratio (CLR) transformation, with principal component analysis used to visualize overall composition as has been described previously (Bennett et al., 2025).

For all analyses, outliers, defined as values exceeding ±2 standard deviations from the group mean, were excluded. All values are reported as mean ± SEM and significance was set at p<0.05. Sample sizes reported in electrophysiology experiments reflect the number of cells recorded and number subjects (cells/n). Data were visualized using GraphPad Prism 11 except for Sankey plots used to describe within-cell transitions in firing phenotype across paired recording conditions, which were generated in MATLAB using the SSankey class (Zhaoxu Liu / slandarer, 2026; sankey plot, MATLAB Central File Exchange).

## Results

### Withdrawal reduces evoked firing in a sex-dependent manner

To determine the effect of acute withdrawal from chronic ethanol exposure on LHb neuron intrinsic excitability, whole-cell patch-clamp recordings were performed 24 h after the final vapor exposure session across the rostro-caudal extent of the LHb (**Fig. 1A**). A multifactorial ANOVA using sex, vapor exposure, and current step as factors was performed to determine the effect of withdrawal on evoked firing in response to positive current injection. This revealed significant main effects of current step [F(10, 605) = 9.850, p < 0.0001] and vapor exposure [F(1, 605) = 40.60, p < 0.0001], as well as a significant sex by vapor interaction [F(1, 605) = 6.669, p = 0.0100], in the absence of a significant main effect of sex [F(1, 605) = 0.6317, p = 0.4270] or three-way interaction [F(10, 605) = 0.8857, p = 0.5464]. The significant sex by group interaction indicated that the magnitude, though not the current-step-dependent shape, of the CIE-induced reduction in evoked firing differed between sexes, justifying separate two-way ANOVAs (group x current step) within each sex to decompose this interaction. In males, this analysis revealed a significant main effect of group [F(1, 297) = 8.348, p = 0.0041] in the absence of a group by current step interaction [F(10, 297) = 0.5569, p = 0.8484], with WD males firing fewer evoked action potentials than AIR males (predicted means: AIR = 5.663, WD = 4.318). In females, a significant main effect of group was similarly observed [F(1, 308) = 35.28, p < 0.0001], again without a significant interaction [F(10, 308) = 0.8912, p = 0.5417], with WD females showing a larger reduction in evoked firing relative to AIR females (predicted means: AIR = 6.298, WD = 3.120) (**Fig. 1B-C**). Together, these data indicate that withdrawal suppresses evoked excitability in the LHb of both sexes, with a significantly greater magnitude of suppression in females.

**Figure 1.**
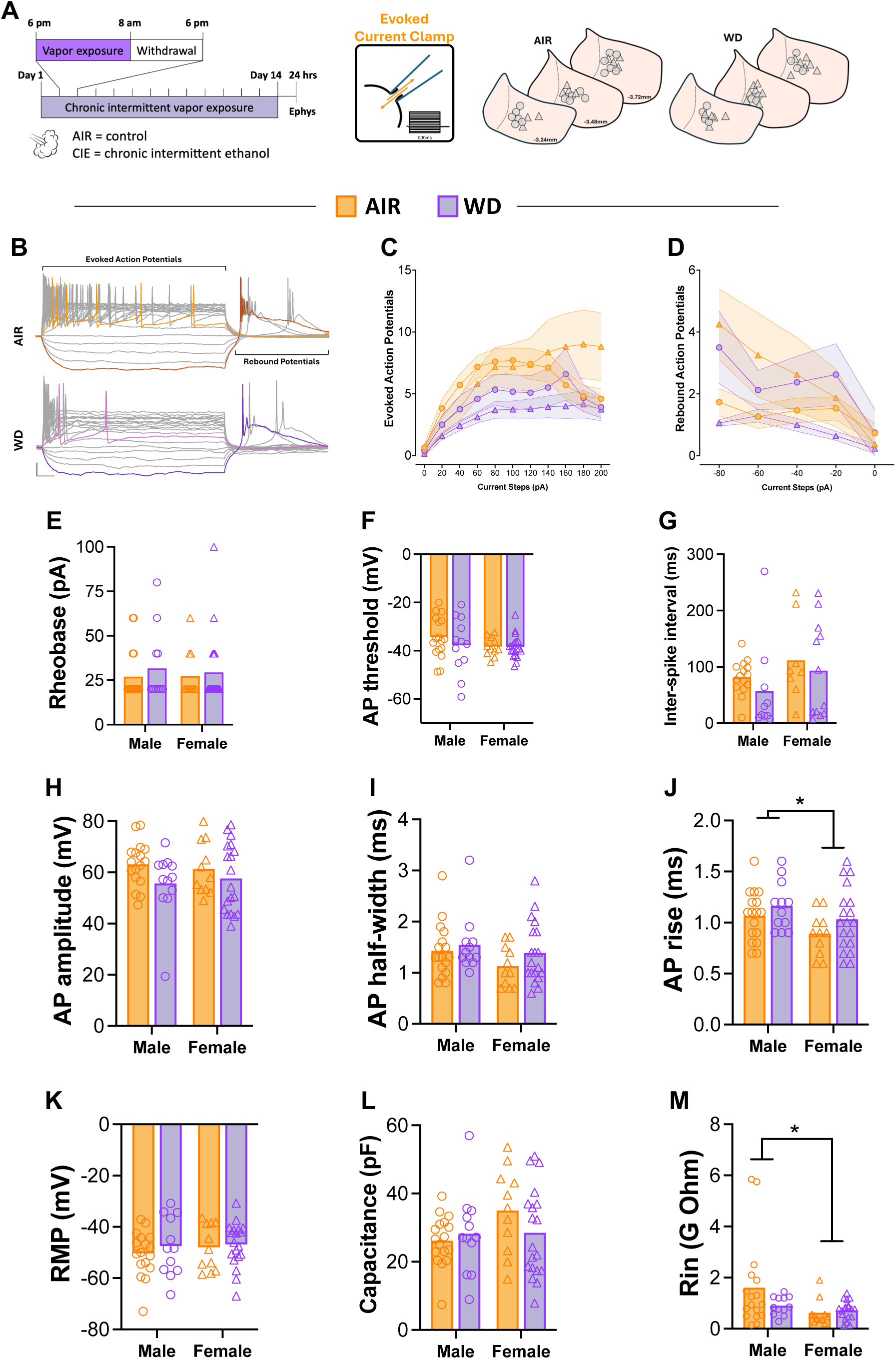
Intrinsic excitability is reduced in LHb neurons of male and female rats during acute withdrawal from chronic ethanol exposure.

Rebound firing following hyperpolarization is a hallmark feature of LHb neurons that has been directly implicated in depression- and anhedonia-related pathology, since T-type calcium-channel-mediated rebound bursts are amplified in models of learned helplessness and are sufficient to drive depressive-like behavior when optogenetically induced (Cui et al., 2018; Y. Yang et al., 2018). We therefore next examined whether withdrawal alters this rebound property in LHb neurons. This revealed a significant main effect of current step [F(4, 220) = 7.570, p < 0.0001] and a significant sex by vapor interaction [F(1, 220) = 26.17, p < 0.0001], in the absence of significant main effects of sex [F(1, 220) = 0.3595, p = 0.5494] or vapor [F(1, 220) = 1.960, p = 0.1629], or a three-way interaction, which approached, but did not reach significance [F(4, 220) = 2.333, p = 0.0567] (**Fig. 1D**). The absence of a significant main effect of group alongside a highly significant sex by group interaction indicates a crossover interaction, in which opposing effects of CIE exposure in males and females cancel when collapsed across sex; this effect would not have been detected using a sex-collapsed analysis alone. Sex-separated two-way ANOVAs performed to decompose this interaction revealed significant main effects of group in both males [F(1, 105) = 6.173, p = 0.0145] and females [F(1, 115) = 23.75, p < 0.0001], again without significant group by current step interactions in either sex (males: F(4, 105) = 0.5608, p = 0.6916; females: F(4, 115) = 2.207, p = 0.0725). Notably, the direction of this effect diverged by sex: WD males exhibited significantly more rebound action potentials than AIR males (predicted means: AIR = 1.347, WD = 2.275), whereas WD females exhibited significantly fewer rebound action potentials than AIR females (predicted means: AIR = 2.475, WD = 0.8471). Together, these findings indicate that acute withdrawal produces a sex-divergent, opposing adaptation in post-inhibitory rebound firing, increasing rebound excitability in males while suppressing it in females, a pattern of genuine directional divergence rather than a difference in effect magnitude alone.

To characterize the intrinsic and action potential (AP) properties underlying these changes, rheobase was identified for each cell and the corresponding trace was used to measure ISI as well as AP threshold, amplitude, rise time, and half-width from the first evoked spike. Two-way ANOVAs revealed no significant main effects or interaction for rheobase (interaction: F(1, 55) = 0.06625, p = 0.7978; sex: F(1, 55) = 0.04480, p = 0.8332; group: F(1, 55) = 0.5302, p = 0.4696; **Fig. 1E**), AP threshold (interaction: F(1, 55) = 0.5857, p = 0.4474; sex: F(1, 55) = 1.321, p = 0.2554; group: F(1, 55) = 0.7356, p = 0.3948; **Fig. 1F**), ISI (no significant differences; **Fig. 1G**), AP amplitude (interaction: F(1, 55) = 0.3759, p = 0.5423; sex: F(1, 55) = 0.0007, p = 0.9791; group: F(1, 55) = 3.302, p = 0.0746; **Fig. 1H**), or AP half-width (interaction: F(1, 55) = 0.2501, p = 0.6190; sex: F(1, 55) = 2.524, p = 0.1179; group: F(1, 55) = 1.629, p = 0.2073; **Fig. 1I**), indicating that these core measures of spike generation were not durably altered by CIE exposure during acute withdrawal. A significant main effect of sex was, however, observed for AP rise time [F(1, 55) = 4.677, p = 0.0349], with females exhibiting significantly faster AP rise times than males independent of vapor exposure, in the absence of a significant effect of group [F(1, 55) = 2.842, p = 0.0975] or interaction [F(1, 55) = 0.1146, p = 0.7363] (**Fig. 1J**).

Passive membrane properties, including resting membrane potential (interaction: F(1, 55) = 0.1251, p = 0.7250; sex: F(1, 55) = 0.3781, p = 0.5412; group: F(1, 55) = 0.6110, p = 0.4377; **Fig. 1K**) and capacitance (interaction: F(1, 55) = 1.975, p = 0.1656; sex: F(1, 55) = 2.221, p = 0.1419; group: F(1, 55) = 0.5445, p = 0.4637; **Fig. 1L**), did not differ by group or sex. A significant main effect of sex was observed for input resistance (Rin) [F(1, 55) = 4.949, p = 0.0302], with females exhibiting significantly lower Rin than males regardless of exposure condition (predicted means: male = 1.258 GΩ, female = 0.6739 GΩ), in the absence of a significant effect of group [F(1, 55) = 1.308, p = 0.2576] or interaction [F(1, 55) = 2.369, p = 0.1295] (**Fig. 1M**). Together, these findings suggest that while gross measures of single-spike AP kinetics are largely unaffected by acute withdrawal, baseline sex differences exist in LHb neuron AP rise time and membrane resistance that are independent of ethanol exposure.

### Spontaneous synaptic transmission on LHb neurons is unaffected by withdrawal

To determine whether CIE exposure produces lasting changes in synaptic drive onto LHb neurons, sEPSCs and sIPSCs were recorded from the same LHb neurons (**Fig. 2A**). Two-way ANOVAs revealed no significant main effects of either vapor or sex or an interaction for sEPSC frequency (interaction: F(1, 57) = 2.161, p = 0.1470; sex: F(1, 57) = 0.6459, p = 0.4249; group: F(1, 57) = 0.0060, p = 0.9384; **Fig. 2B-C**) or amplitude (interaction: F(1, 57) = 0.7791, p = 0.3811; sex: F(1, 57) = 0.1166, p = 0.7340; group: F(1, 57) = 0.00009, p = 0.9923; **Fig. 2D**). Analysis of sIPSC frequency revealed a significant main effect of sex [F(1, 29) = 8.075, p = 0.0081], with no significant effect of vapor [F(1, 29) = 0.0060, p = 0.9386] or interaction [F(1, 29) = 1.186, p = 0.2850] (**Fig. 2E-F**). No significant effects of vapor or sex were observed for sIPSC amplitude (interaction: F(1, 29) = 0.4808, p = 0.4936; sex: F(1, 29) = 1.784, p = 0.1921; group: F(1, 29) = 3.378, p = 0.0763; **Fig. 2G**). Analysis of the balance of excitatory and inhibitory transmission using E/I frequency ratio (interaction: F(1, 32) = 0.0028, p = 0.9580; sex: F(1, 32) = 0.4173, p = 0.5229; group: F(1, 32) = 0.0656, p = 0.7996; **Fig. 2H-I**) and synaptic drive (interaction: F(1, 32) = 0.1189, p = 0.7325; sex: F(1, 32) = 0.0341, p = 0.8547; group: F(1, 32) = 0.3691, p = 0.5478; **Fig. 2J**) revealed no significant differences between vapor groups or sexes. Altogether, these data indicate that spontaneous excitatory and inhibitory postsynaptic drive onto LHb neurons is not altered during acute withdrawal from chronic ethanol exposure.

**Figure 2.**
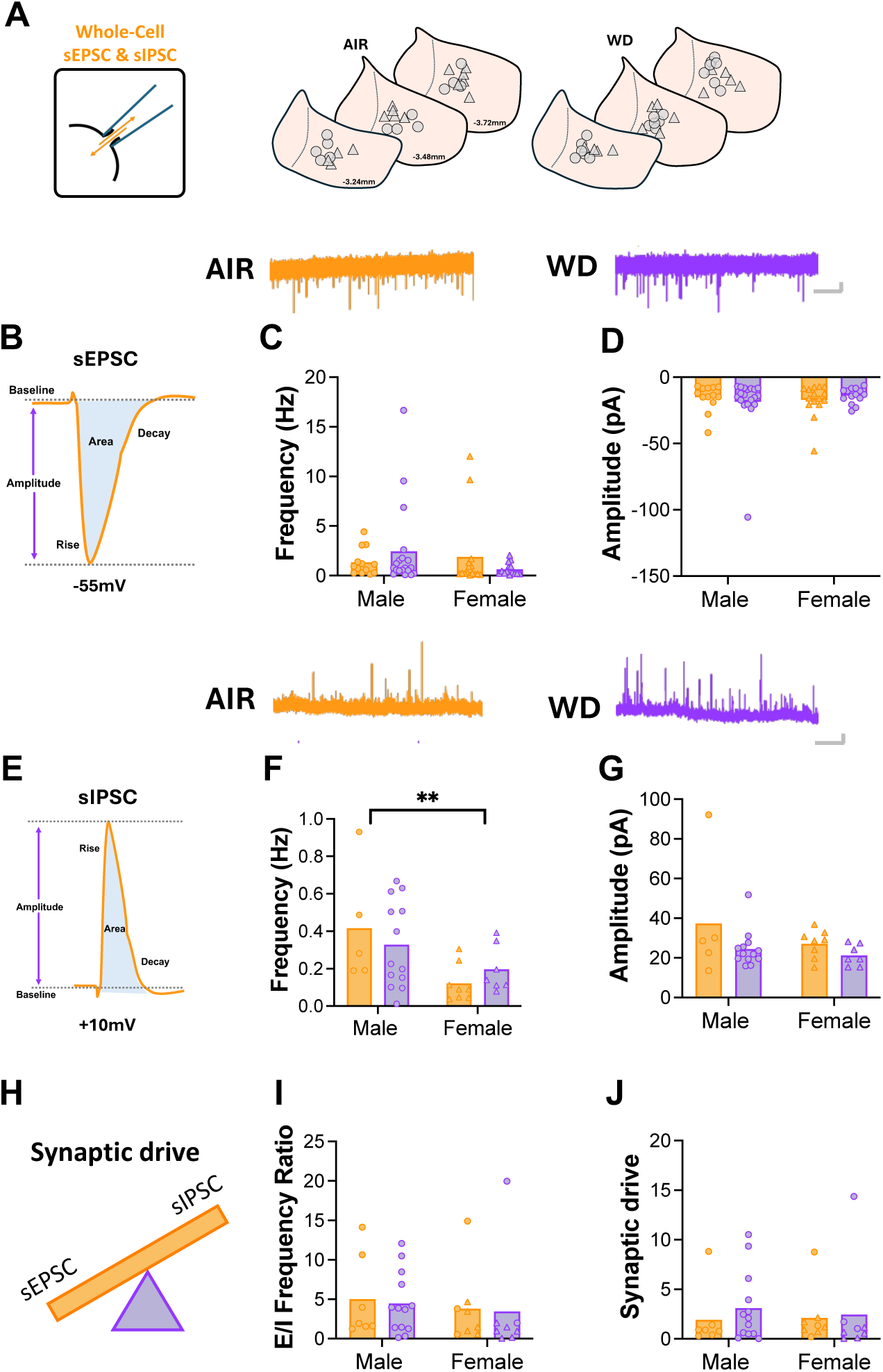
Spontaneous synaptic transmission is unaltered in the LHb during acute withdrawal from chronic ethanol exposure.

### Withdrawal reduces spontaneous LHb firing without altering firing phenotype dynamics

Cell-attached recordings were used to assess the effect of withdrawal on the spontaneous, unperturbed firing activity of LHb neurons (**Fig. 3A**). Similar to the effect of withdrawal on evoked firing, a two-way ANOVA of mean spontaneous firing frequency revealed a significant main effect of vapor [F(1, 107) = 6.002, p = 0.0159], with WD rats exhibiting significantly lower spontaneous firing frequency relative to AIR controls (predicted means: AIR = 5.504 Hz, WD = 3.015 Hz). This occurred in the absence of a significant effect of sex [F(1, 107) = 0.8810, p = 0.3500] or vapor by sex interaction [F(1, 107) = 0.1480, p = 0.7012] (**Fig. 3B**). No significant effects of vapor or sex were observed for ISI (interaction: F(1, 107) = 0.01603, p = 0.8995; sex: F(1, 107) = 0.5649, p = 0.4539; group: F(1, 107) = 2.057, p = 0.1544; **Fig. 3C**).

**Figure 3.**
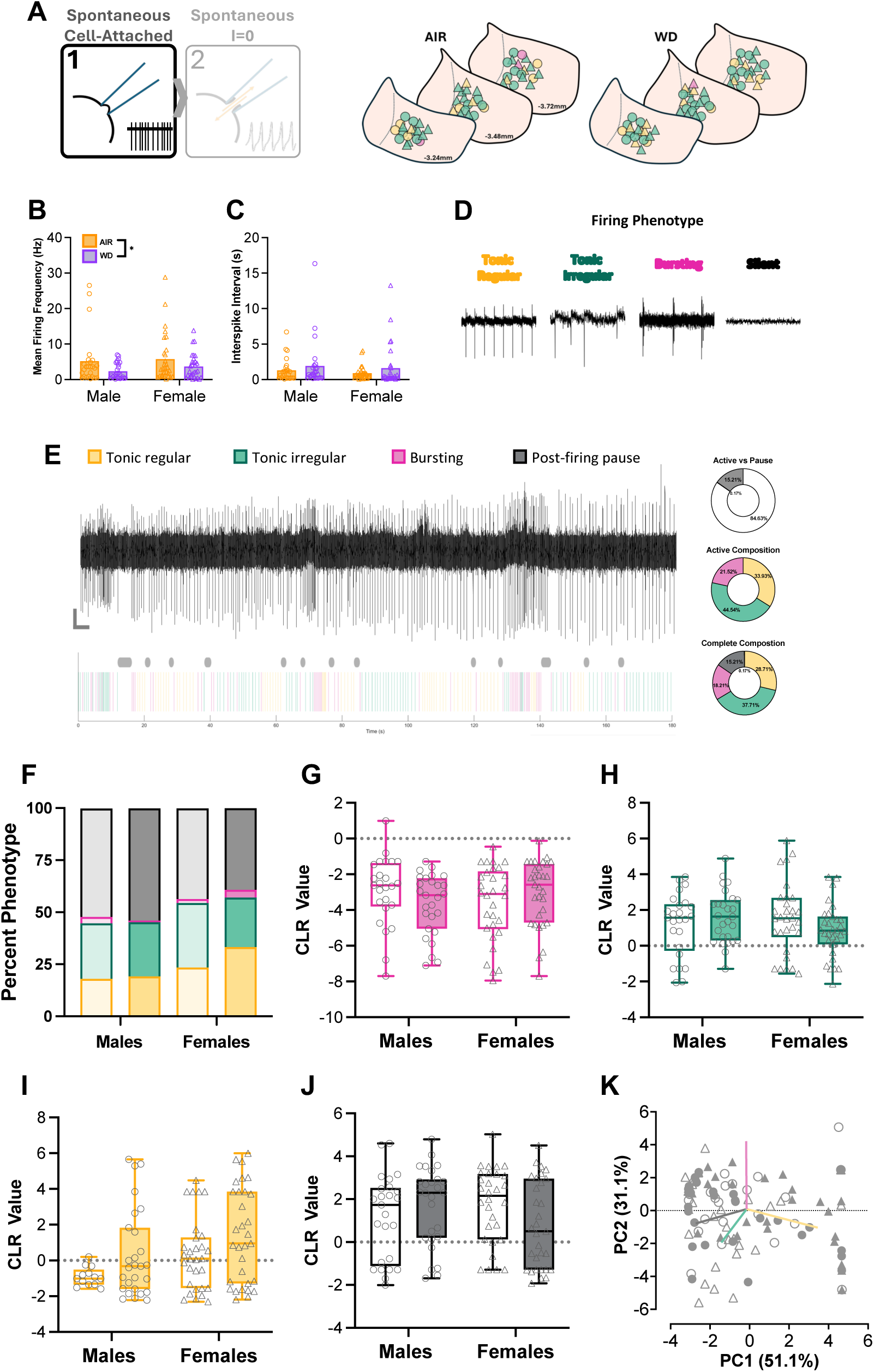
Spontaneous firing is reduced during acute withdrawal in the absence of changes to stochastic firing patterns.

To determine whether withdrawal altered the balance among distinct LHb firing patterns, we next classified activity segments during each recording as tonic regular, tonic irregular, or burst, with periods of silence between activity segments classified as a post-firing pause (**Fig. 3D**). **Fig. 3E** shows a representative 3-min cell-attached recording and MATLAB-based phenotype classification for a representative LHb neuron. When the proportion of each phenotype was compared across group and sex (**Fig. 3F**), no evident shift in overall phenotype composition was apparent between AIR and WD rats of either sex. Compositional log-ratio (CLR) analysis of burst (**Fig. 3G**), tonic irregular (**Fig. 3H**), tonic regular (**Fig. 3I**), and pause (**Fig. 3J**) phenotypes did not reveal significant effects of either vapor or sex. Consistent with this, a PCA biplot incorporating all phenotype CLR values did not reveal separation of WD from AIR cells by either vapor group or sex (**Fig. 3K**). Together, these findings indicate that while CIE exposure reduces the overall rate of spontaneous LHb firing during early withdrawal, the underlying firing phenotype composition of these neurons remains stable.

### LHb firing phenotype is dynamic across recording configurations independent of withdrawal

To determine whether the firing phenotype of LHb neurons is preserved across recording configurations, a subset of cells that were recorded in cell-attached configuration were subsequently recorded in whole-cell configuration (**Fig. 4A**). Representative 3-min I=0 recordings with paired MATLAB-based phenotype quantification are shown in **Fig. 4B**. As in cell-attached recordings, the proportion of each phenotype did not differ appreciably by vapor or sex (**Fig. 4C**). CLR analysis did not reveal significant effects of vapor or sex for burst (**Fig. 4D**), tonic irregular (**Fig. 4E**), tonic regular (**Fig. 4F**), or pause (**Fig. 4G**) phenotypes, and a PCA biplot of these values did not reveal separation by group or sex (**Fig. 4H**). To assess the stability of firing phenotype classification across recording configurations, we compared each cell’s phenotype in the cell-attached configuration to its classification following break-in (I=0) (**Fig. 4I**). Phenotype classification was maintained in less than half of all cells overall (27/61, 44.3%). Although concordance appeared numerically higher in female AIR (10/19, 52.6%) and male WD (9/16, 56.3%) cells compared to female WD (6/17, 35.3%) and male AIR (2/9, 22.2%) cells, this difference did not reach statistical significance. These data suggest that firing phenotype classified by cell-attached recording is not consistently predictive of whole-cell firing phenotype. Together, these findings indicate that individual LHb neurons dynamically fluctuate across firing phenotypes rather than possessing a single, fixed identity. Moreover, acute withdrawal does not alter the prevalence of any given phenotype. The reduction in firing rate observed in WD animals (**Fig. 3B**) therefore likely reflects a change in overall output rate rather than a shift in firing pattern

**Figure 4.**
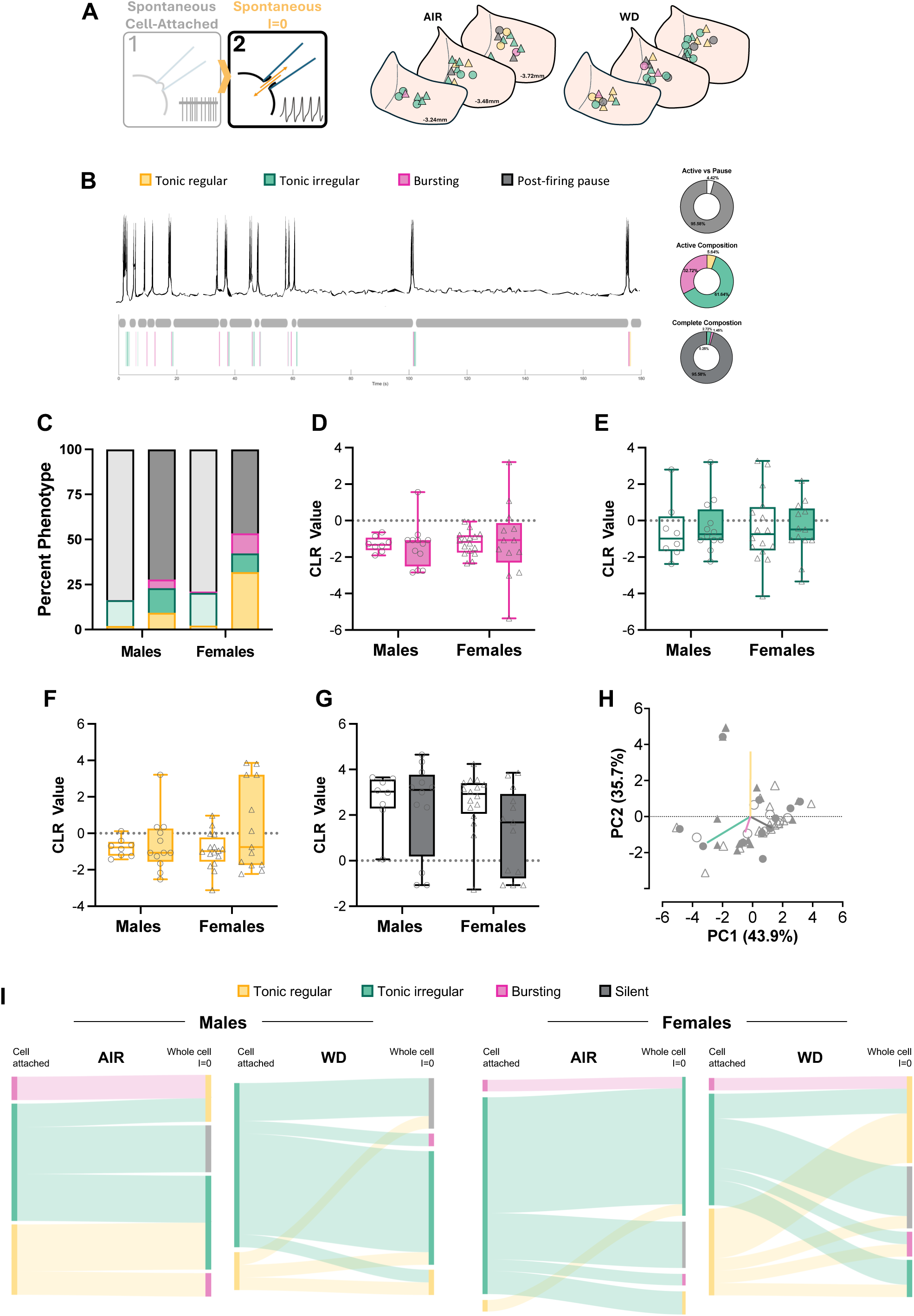
Spontaneous firing phenotype in not maintained across recording configurations.

### Withdrawal-induced increase in astrocyte density is associated with disrupted astrocytic regulation of synaptic glutamate

Given the reduction in spontaneous LHb firing observed during withdrawal in the absence of changes in fast synaptic transmission or firing phenotype, we next considered alternative mechanisms by which withdrawal may alter excitability. Astrocytes actively regulate synaptic transmission through perisynaptic processes that form ‘tripartite synapses’ with neurons, sensing neurotransmitter release and shaping synaptic signaling via glutamate transporters like GLT-1 (Araque et al., 1999; Perea et al., 2009). This astrocytic compartment is also a well-established target of chronic ethanol exposure, which produces long-lasting changes in astrocyte gene expression, morphology, and proliferation across brain regions (Adermark & Bowers, 2016; Erickson et al., 2018). We therefore asked whether withdrawal produces analogous astrocytic changes in the LHb that could account for the observed reduction in firing. To explore this, we used immunofluorescence staining to measure changes in astrocyte density in the LHb (**Fig. 5A**). An unpaired t-test revealed a significant increase in Sox9 expression in the LHb of WD rats relative to AIR controls [t(21) = 2.175, p = 0.0412] (AIR = 576.4, WD = 683.7; mean difference = 107.3 ± 49.32; **Fig. 5B**). Consistent with this, GFAP expression was also significantly higher in the LHb of WD rats relative to AIR controls [t(21) = 2.208, p = 0.0385] (AIR = 0.05396, WD = 0.09123; mean difference = 0.03727 ± 0.01688; **Fig. 5C**). Together, these data suggest that withdrawal from chronic ethanol exposure increases astrocytic proliferation in the LHb.

**Figure 5.**
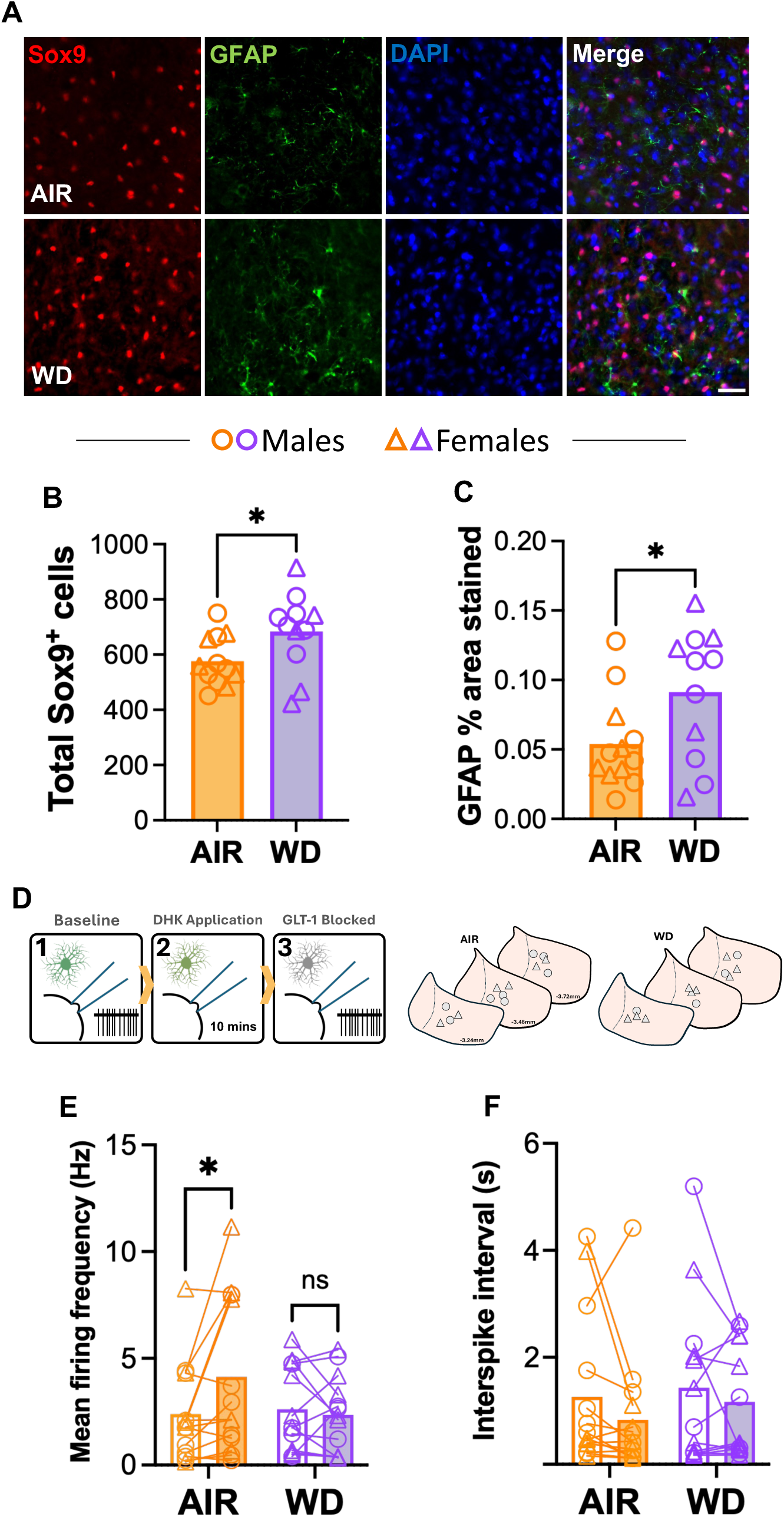
Withdrawal-induced alterations in astrocyte structure and function.

To determine whether this astrocytic hypertrophy was accompanied by changes in astrocyte-dependent regulation of neuronal activity, we measured spontaneous firing in LHb neurons before and during inhibition of the astrocyte-specific glutamate transporter, GLT-1, DHK (**Fig. 5D**). A two-way repeated-measures ANOVA of mean firing frequency revealed a significant vapor by drug interaction [F(1, 25) = 5.768, p = 0.0241] in the absence of significant main effects of either vapor [F(1, 25) = 0.7606, p = 0.3914] or drug [F(1, 25) = 3.103, p = 0.0904]. Šídák’s post hoc comparisons revealed that DHK application significantly increased firing frequency relative to baseline in AIR rats (predicted means: pre-bath = 2.385 Hz, DHK = 4.138 Hz; p = 0.0120) but that this effect was absent in WD rats (predicted means: pre-bath = 2.620 Hz, DHK = 2.351 Hz; p = 0.8847) (**Fig. 5E**). No significant effects of vapor exposure, drug application, or an interaction were observed for ISI (interaction: F(1, 25) = 0.1641, p = 0.6888; vapor: F(1, 25) = 0.3007, p = 0.5883; drug: F(1, 25) = 2.776, p = 0.1082; **Fig. 5F**). Together, these findings indicate that withdrawal from chronic ethanol exposure disrupts astrocytic glutamate transporter-dependent regulation of spontaneous LHb neuron firing.

## Discussion

The present study examined physiological alterations in the LHb during acute withdrawal in a rat model of alcohol dependence. Our findings unexpectedly reveal a significant decrease in both evoked and spontaneous firing in the LHb of withdrawn rats compared to controls. Notably, while this effect was observed in both sexes, the effect of withdrawal to suppress LHb firing was larger in magnitude in females than in males. Interestingly, hyperpolarizing-induced rebound firing was altered in a sex-divergent manner, with males exhibiting more rebound firing, whereas females exhibited less during withdrawal compared to controls. The overall dynamic nature of LHb firing patterns was preserved during withdrawal indicating that chronic ethanol exposure reduces overall firing rate rather than reorganizing the pattern of LHb output. The withdrawal-induced decrease in firing occurred in the absence of alterations in spontaneous excitatory or inhibitory postsynaptic currents or in intrinsic membrane properties. Instead, these changes were accompanied by an increase in astrocyte density and a loss of astrocyte-mediated glutamate clearance. Taken together, our findings describe a withdrawal state in which the astrocytic compartment is structurally reactive yet functionally impaired while neuronal output is reduced pointing to significant disruptions in astrocyte-neuron coupling in the LHb after chronic ethanol exposure.

We found that acute withdrawal reduced both intrinsic excitability and spontaneous firing in LHb neurons, an effect that occurred in the absence of any accompanying change in spontaneous synaptic transmission. These results are in contrast to previous work showing that repeated ethanol exposure increases evoked firing in the LHb (Li et al., 2016; Li et al., 2017; Kang et al., 2019; Gregor et al., 2019; Flanigan et al., 2023; Kang et al., 2017, 2018). However, animals in these studies were exposed to ethanol using either repeated intraperitoneal injection or voluntary drinking paradigms, neither of which reliably produces the sustained blood ethanol concentrations or physical dependence characteristic of vapor-based exposure models. Moreover, even when long-term drinking does produce measurable withdrawal, the severity of withdrawal symptoms is likely to be variable given individual differences in intake that is inherent to volitional drinking unlike the repeated, high-dose intoxication-withdrawal cycles characteristic of CIE vapor exposure (Gilpin et al., 2008, 2009; Glover et al., 2019, 2021). It is therefore possible that the discrepancy between our findings and this prior work reflects differences in the volitional versus passive nature of ethanol exposure across models or in the severity of dependence achieved. Indeed, variability in LHb intrinsic excitability after long-term ethanol drinking observed in previous work lends support for the latter possibility (Gregor et al., 2019; Kang et al., 2019). In addition, much of the aforementioned work performed ethanol exposure during adolescence and the majority of studies were limited to male subjects. The latter methodological difference may be of particular relevance given our observation that withdrawal from CIE exposure produced an even greater reduction in intrinsic excitability in the LHb of females than males.

Our findings are also unexpected considering prior work from our own group and others revealing significant cFos induction in the LHb during acute withdrawal from CIE vapor exposure (Glover et al., 2019; Nentwig et al., 2022). Several factors may reconcile these seemingly discrepant results. First, it is possible that inputs driving spontaneous firing are lost in the *ex vivo* slice preparation and thus obscuring a potential increase in firing that is more accurately captured in cFos experiments reflective of intact afferent connections. Indeed, previous work has shown that loss of input from the entopeduncular nucleus, which is severed in our acute slice preparation, diminishes spontaneous and burst firing in the LHb (H. Li et al., 2019). Considering this, it is possible that while baseline, spontaneous LHb activity is reduced during acute withdrawal, responses evoked by synaptic input or environmental stimuli may be selectively augmented. Such a state-dependent effect would only be detectable using *in vivo* approaches capable of capturing stimulus-driven activity in the intact, behaving animal. Alternatively, or in addition, the reduced intrinsic excitability we observe in the LHb during acute withdrawal may represent a compensatory, homeostatic adaptation that arises in response to, and ultimately offsets, augmented activity occurring elsewhere in the intact network, such as at the level of afferent drive, which cannot be captured using *ex vivo* recordings.

It is also important to consider that, although conventionally used as an indicator of recent neuronal activation, cFos expression is not neuron-specific. In fact, previous work has shown that astrocytes, microglia, and oligodendrocytes all have the ability to express cFos (Aguilar-Delgadillo et al., 2024; Condorelli et al., 1993; Groves et al., 2018; Anderson et al., 1994; Morgan & Curran, 1991). cFos induction in non-neuronal cells is particularly common in response to neuroinflammation – a process consistently observed after chronic ethanol exposure (Lékó et al., 2023). Our observation of increased Sox9 and GFAP expression in the LHb of withdrawn rats relative to controls lends support for the possibility that LHb cFos induction observed in previous studies is due, at least in part, to astrogliosis. Sox9 serves as a robust, astrocyte-specific nuclear marker across most CNS regions, enabling accurate quantification of astrocyte number independent of membrane-associated markers, which are often reflective of hypertrophy (Sun et al., 2017). The increase in Sox9+ cell counts we observe therefore reflects a genuine increase in astrocyte number in the LHb during withdrawal. This interpretation is reinforced by our GFAP findings, as chronic ethanol exposure is known to increase GFAP synthesis and the number and diameter of GFAP-positive astrocytes across white and gray matter (Dalçik et al., 2009). Thus, our findings are consistent with either an increase in astrocyte number, an expansion of individual astrocyte territory, or both.

At first glance, our evidence for astrocyte proliferation and/or hypertrophy appears difficult to reconcile with our finding that DHK-sensitive, GLT-1-mediated glutamate clearance is functionally impaired in the LHb during acute withdrawal. However, this apparent paradox is not inconsistent with the broader literature, as the relationship between astrocyte density or structural markers and glutamate transporter function is not strictly one-to-one. For example, a number of studies have shown increased astrocytic markers in tandem with reduced astrocyte function (Campbell et al., 2020; Schreiner et al., 2013). This dissociation extends to the transporter itself, as reactive astrocytes in a mouse model of Alzheimer’s disease showed impaired glutamate transporter currents despite unchanged GLT-1 protein levels (Srivastava et al., 2025). Conversely, measures of astrocytic dysfunction are not always associated with changes in density or morphology (Schmitz et al., 2016), possibly reflecting changes in transporter trafficking or localization that are not captured by bulk expression measures. Glutamate clearance instead depends on the physical relationship between astrocytic processes and individual synapses, with smaller spines receiving proportionally greater astrocytic coverage and more efficient glutamate uptake than larger ones (Herde et al., 2020). By this logic, the observed increase in astrocyte density in the withdrawn LHb does not guarantee a proportional increase in functional glutamate clearance and instead may reflect a population of astrocytes that is structurally reactive but functionally compromised.

The loss of GLT-1-mediated glutamate clearance we observed in the LHb during acute withdrawal could translate to increased extrasynaptic glutamate levels, which may in turn serve as a mechanism for increased excitation of LHb neurons *in vivo*, an effect that could go undetected in the slice preparation and would not necessarily be captured by measures of discrete synaptic currents, since tonic extrasynaptic glutamate acts independently of the frequency or amplitude of spontaneous synaptic events (Fleming et al., 2011; Sah et al., 1989). This interpretation is consistent with the established capacity of neuromodulators and ambient extracellular transmitter to adjust neuronal gain independently of discrete synaptic weight (Marder, 2012; Nadim & Bucher, 2014). It is possible that this same mechanism contributes to a homeostatic reduction in intrinsic excitability, of the kind we observe in our evoked firing experiments, as LHb neurons compensate for chronically elevated extrasynaptic glutamate tone. This is further supported by evidence that the LHb possesses its own activity-dependent homeostatic machinery: acute stress engages, while chronic stress suppresses, an autophagy-based mechanism that regulates LHb excitability and synaptic transmission through on-demand degradation of glutamate receptors to defend a set-point of activity (L. Yang et al., 2025), demonstrating that the LHb is capable of exactly this kind of compensatory, activity-dependent recalibration.

In addition to reduced evoked and spontaneous firing, withdrawal produced a sex-divergent change in rebound firing following hyperpolarization, increasing it in males while decreasing it in females. In the LHb, membrane hyperpolarization is a well-established trigger for rebound burst firing, with brief hyperpolarizing input capable of driving long-lasting discharges of action potentials in the majority of LHb neurons (Chang & Kim, 2004). This rebound firing depends on T-type calcium channels acting together with NMDA receptors. Bursting through this same mechanism has been directly linked to depressive-like behavior with blockade of either channel in the LHb producing rapid antidepressant-like effects (Y. Yang et al., 2018). Interestingly, astrocytes gate this excitability in the LHb via astroglial Kir4.1, which controls extracellular potassium levels thereby regulating the degree of hyperpolarization and resulting burst activity (Cui et al., 2018). This raises the possibility that the astrocytic changes we observe during acute withdrawal shape not only overall firing rate but also this sex-divergent rebound phenotype. Because rebound firing determines how strongly the LHb responds following a period of inhibition, the sex differences we observe suggest that males and females respond to inhibitory input with opposing effects on net excitability. Importantly, this is distinct from the general withdrawal-induced suppression of activity described above indicating that this adaptation is not simply an extension of the observed changes in evoked-firing but a distinct, sex-dependent process. Whether this divergence in rebound excitability translates into sex differences in withdrawal-related behavior is an important consideration for future experiments.

Prior work has characterized LHb neurons into distinct firing phenotypes, including tonic regular, tonic irregular, bursting, and silent, with data suggesting that these phenotypes are stable and tied to intrinsic membrane properties (Wagner et al., 2017; Weiss & Veh, 2011). In contrast, firing phenotype in our recordings was stochastic and dynamic and was not consistently maintained within the same cell across minutes of recording or across recording configurations. This suggests that firing phenotype, classified at a single moment or under a single recording condition, is unlikely to reflect a stable trait-like property of an individual LHb neuron. Rather, it likely reflects a state that fluctuates depending on the immediate physiological context of the cell. Interestingly, the relative proportion of tonic, burst, and pause activity did not differ by vapor exposure or sex, despite the fact that mean firing frequency was reduced during withdrawal. This is suggestive of a uniform downward scaling of neuronal gain rather than a reorganization of firing phenotypes and is consistent with homeostatic plasticity mechanisms that uniformly scale neuronal output or excitability in response to activity, rather than selectively reorganizing individual synapses or firing patterns (Desai et al., 1999; Turrigiano, 2008; van Rossum et al., 2000).

In conclusion, our findings identify the LHb as a site of coordinated neuronal and astrocytic adaptation during acute withdrawal from chronic ethanol exposure. Rather than the straightforward hyperexcitability predicted by prior models of ethanol-induced LHb dysfunction, our data reveal a more complex picture in which reduced neuronal output coexists with structurally reactive but functionally impaired astrocytes, and in which sex shapes not only the magnitude but the direction of specific physiological adaptations. Taken together, our results underscore the need for future work investigating the effects of ethanol exposure on the LHb by pairing *in vivo* and *ex vivo* recordings to better link changes in astrocyte function with neuronal output.

## Acknowledgements

This work was supported by NIH grants R01 AA031003, R01 AA029130, and P50 AA922537 to EJG and T32 AA0236577 to KYC.

